# A deep learning approach to real-time HIV outbreak detection using genetic data

**DOI:** 10.1101/2021.12.17.473204

**Authors:** Michael D. Kupperman, Thomas Leitner, Ruian Ke

## Abstract

Pathogen genomic sequence data are increasingly made available for epidemiological monitoring. A main interest is to identify and assess the potential of infectious disease outbreaks. While popular methods to analyze sequence data often involve phylogenetic tree inference, they are vulnerable to errors from recombination and impose a high computational cost, making it difficult to obtain real-time results when the number of sequences is in or above the thousands.

Here, we propose an alternative strategy to outbreak detection using genomic data based on deep learning methods developed for image classification. The key idea is to use a pairwise genetic distance matrix calculated from viral sequences as an image, and develop convolutional neutral network (CNN) models to classify areas of the images that show signatures of active outbreak, leading to identification of subsets of sequences taken from an active outbreak. We showed that our method is efficient in finding HIV-1 outbreaks with *R*_0_ *≥* 3, and overall a specificity exceeding 85% and sensitivity better than 70%. We validated our approach using data from HIV-1 CRF01 in Europe, containing both endemic sequences and a well-known dual outbreak in intravenous drug users. Our model accurately identified known outbreak sequences in the background of slower spreading HIV. Importantly, we detected both outbreaks early on, before they were over, implying that had this method been applied in real-time as data became available, one would have been able to intervene and possibly prevent the extent of these outbreaks. This approach is scalable to processing hundreds of thousands of sequences, making it useful for current and future real-time epidemiological investigations, including public health monitoring using large databases and especially for rapid outbreak identification.

**Author summary:** The analysis of pathogen genomic data to analyze epidemics at scale is constrained by the computational cost associated with phylogenetic tree reconstruction. As a fast and efficient alternative, we employed convolutional neural networks to analyze evolutionary pairwise distance matrices as images to perform classifications of the current epidemiological situation of a growing public health sequence database. We used simulated data to train and test our model, and as validation we accurately mapped the start and end of two linked well-documented HIV-1 outbreaks in the backdrop of ongoing slower HIV spread. Thus, our new approach is efficient, accurate, scalable, and can analyze data in real time.

## Introduction

The human population is increasingly exposed to threats of infectious disease outbreaks due to population growth, increased frequency of traveling, changing patterns of land use etc. This is exemplified by the ever-growing HIV epidemic [1] as well as recent emerging outbreaks such as the SARS-CoV-2 pandemic. A key to outbreak control is early detection when the number of infected individuals is small and the disease spread is local. One growing resource for disease control is to utilize pathogen genomic sequence data to assess epidemiological conditions and threats.

Given sequence data, state-of-the-art, flexible phylogenetic methods have been developed for analysis of general evolutionary questions [2–4], applicable to pathogen evolution, as well as faster but less precise algorithms for large data [5]. More focused phylodynamic methods have also been developed for specific tasks, e.g., taking incidence data into account [6], including multiple evolutionary scales [7], inferring underlying transmission networks on several levels [8, 9], and using large next generation sequencing data [10]. Motivated by pathogen evolution, advanced methods for inference of past demographic history with population size dynamics and migration that can reconstruct outbreaks have also been developed, e.g., the modular framework of BEAST [11], which takes a Bayesian approach to account for uncertainties in the tree reconstruction. However, phylogenetic tree reconstruction, interpretation, and subsequent outbreak identification requires extensive expert knowledge, and thus typically can be reliably done only by highly trained scientists. Furthermore, despite the advances and promises of genomic epidemiology, current phylogenetic methods to analyze pathogen genetic data suffer from high computational cost that prevents real-time or near real-time analysis, especially when thousands of sequences need to be analyzed, as is common in modern sequencing projects.

Here, we propose an alternative to this paradigm. Instead of constructing phylogenetic trees, we based our approach on the pairwise distance matrix of a sample of genetic sequences, and used a deep learning approach to analyze the matrix. Deep learning approaches, such as the convolutional neural network (CNN) [12], has been well developed and widely used for image identification over the past decade [13–15]. By using many parameters in a highly nonlinear model, a deep learning model can efficiently learn complex relationships within the data to form highly accurate predictions [16]. Our rationale here is that outbreaks would lead to distinctive signatures in the pairwise distance matrix (as well as impact the topology of the phylogenetic tree [17]). We leveraged the advance in deep learning models by treating the matrix as an image, and developed deep learning models to identify these signatures from the pairwise distance matrix and thus detect outbreaks from a sequence database. We show that our CNN models made accurate predictions against historical HIV sequence data with known epidemiological history, and that they can handle many thousands of sequences within a very short time frame using a laptop computer.

## Methods

### Overview of the framework

Here we describe the main workflow of our approach. We first developed a forward stochastic simulator of HIV transmission to generate synthetic datasets for training and testing of our CNN models. In this simulator, the number of infected individuals initially expand exponentially, and subsequently establish a constant population size (Fig. 1A). The entire transmission history was recorded; *n* individual samples (*n* = 15, 20, 30, 40 or 50 in our model) were taken either during the exponentially increasing phase (labeled as ‘epidemic’) or the constant population phase (labeled as ‘endemic’). Nucleotide substitutions were then simulated on the transmission tree/genealogy of the *n* sampled individuals, and a *n* × *n* pairwise distance matrix was derived from the simulated substitutions on the transmission tree (Fig. 1B). We repeated the stochastic simulation many times to derive a rich synthetic dataset (i.e. a collection of matrices with labels).

**Fig 1.**
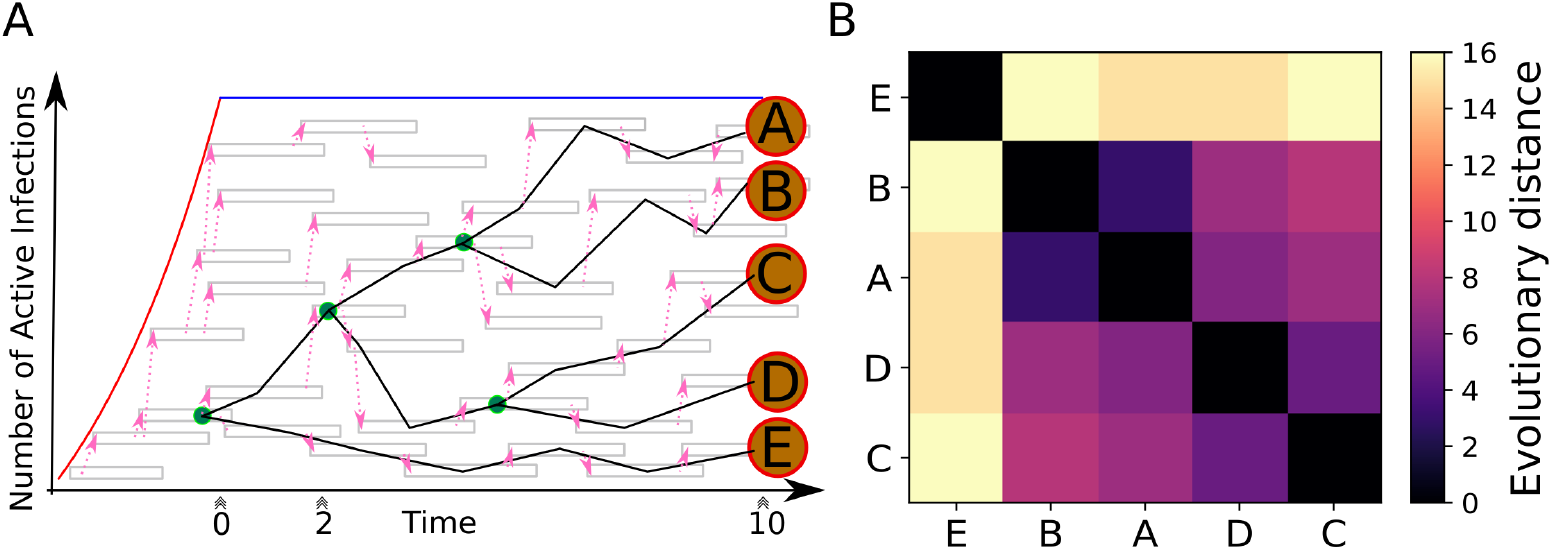
(A) Cartoon representation of the biphasic forwards stochastic simulation. The red line indicates the exponential phase, the blue line indicates the constant population phase. Grey boxes represent an active infection, pink arrows indicate transmission. The reconstructed transmission dendrogram is overlaid, green nodes are internal, orange (lettered) nodes indicate sampled infections. Triple chevrons indicate different possible sampling times (0, 2, 10 yrs). (B) The resulting, sorted pairwise distance matrix image of the sample at 10 years.

We then developed CNN models, trained and tested these CNN models using the synthetic dataset to identify matrices that are labeled as ‘epidemic’. These CNN models handle a small number of sequences (*n* = 15, 20, 30, 40 or 50) at a time ; however, a pathogen database may contain tens of thousands or even hundreds of thousands of sequences. We thus developed a ’sliding-window’ approach to utilize the CNN model to identify subsets of sequences that belong to an active outbreak, i.e. the ‘epidemic’ label, from a collection of a large number of sequences. More specifically, we first constructed a pairwise distance matrix for all the sequences in the database (*m* sequences in total), and reordered the matrix using a fast clustering algorithm, such as hierarchical clustering [18]. This ensures that the sequences belonging to an active outbreak are grouped together. We then used a window of size *n* × *n*, and slide this window from one diagonal of the *m* × *m* matrix towards the other diagonal. At each position of the sliding window, we used the trained CNN model to predict the label (‘epidemic’ or ‘endemic’) for the *n* × *n* submatrix of the sliding window. This sliding window approach is similar to the well-established image identification algorithm, such as R-CNN, where a subarea/window of the entire image is sampled to identify objects of interests [19]. This approach should allow the analysis of tens of thousands of sequences very efficiently.

### The HIV-1 stochastic transmission simulator

The HIV-1 stochastic forward transmission model (Fig 1) was adapted from [20]. It has two phases: 1) an exponential growth phase and, 2) a constant population phase (Fig 1 A). The simulation began with one infected individual. Secondary infections were generated stochastically according to a predefined basic reproductive number *R*_0_(ranging between 1.5 and 5 in our simulations). It has been shown that the transmission potential is much higher during the acute infection phase in an infected individual [21, 22]. Thus, we assumed that during the first three months the rate of new infections is 20-fold higher than the remaining infectious period. We assumed that an individual will be non-infectious when on successful antiviral treatment, administered after diagnosis between 13 to 36 months after infection [23, 24].

Once the population size reaches the maximum number of infected individuals, assumed to be log-uniformly distributed between 10^3^ and 10^4^ across different simulation runs, we set the population size to be constant over time. In the constant population phase, the simulation switched from sampling the number of new infections to only replacing infections that have reached the end of their active phase, maintaining the total number of active infections (Fig 1 A).

The transmission history of all individuals during the simulation was recorded, which allows for the reconstruction of the transmission history/tree for the entire population or any subset thereof. Samples (with a size between 15 and 50) were taken at 3 different time points. The first time point, at 0 years, is when the population size reaches the maximum number of infected individuals, i.e., the transition time when the population changes from exponential growth to a constant size. The genealogical relationship between the samples taken at this time point reflects populations undergoing exponential growth, and therefore, we labeled the samples as ‘epidemic’. The 2nd and 3rd time points are 2 and 10 years after the first time point. The genealogical relationship between these samples therefore reflects populations that have stopped expanding, and we labelled them as ‘endemic’.

#### Construction of Pairwise Distance Matrices

To construct an evolutionary pairwise distance matrix, *D*, we first calculated the temporal distance between each pair of samples based on the transmission history/tree. We assumed that the genetic sequence data is approximately 300 nucleotides (nt) to be consistent with the real HIV-1 data we used below. We then calculated the pairwise genetic distance separating two samples by drawing the number of substitutions from a Poisson distribution with the expected number of substitutions as the mean parameter, here 0.002 substitutions nt^−1^ year^−1^ times 300 nt, times the temporal distance. This was iterated for each pair of the samples to form the pairwise distance matrix. Examples of matrices sampled from each time point (0, 2, or 10 years) are shown in figure 2.

**Fig 2.**
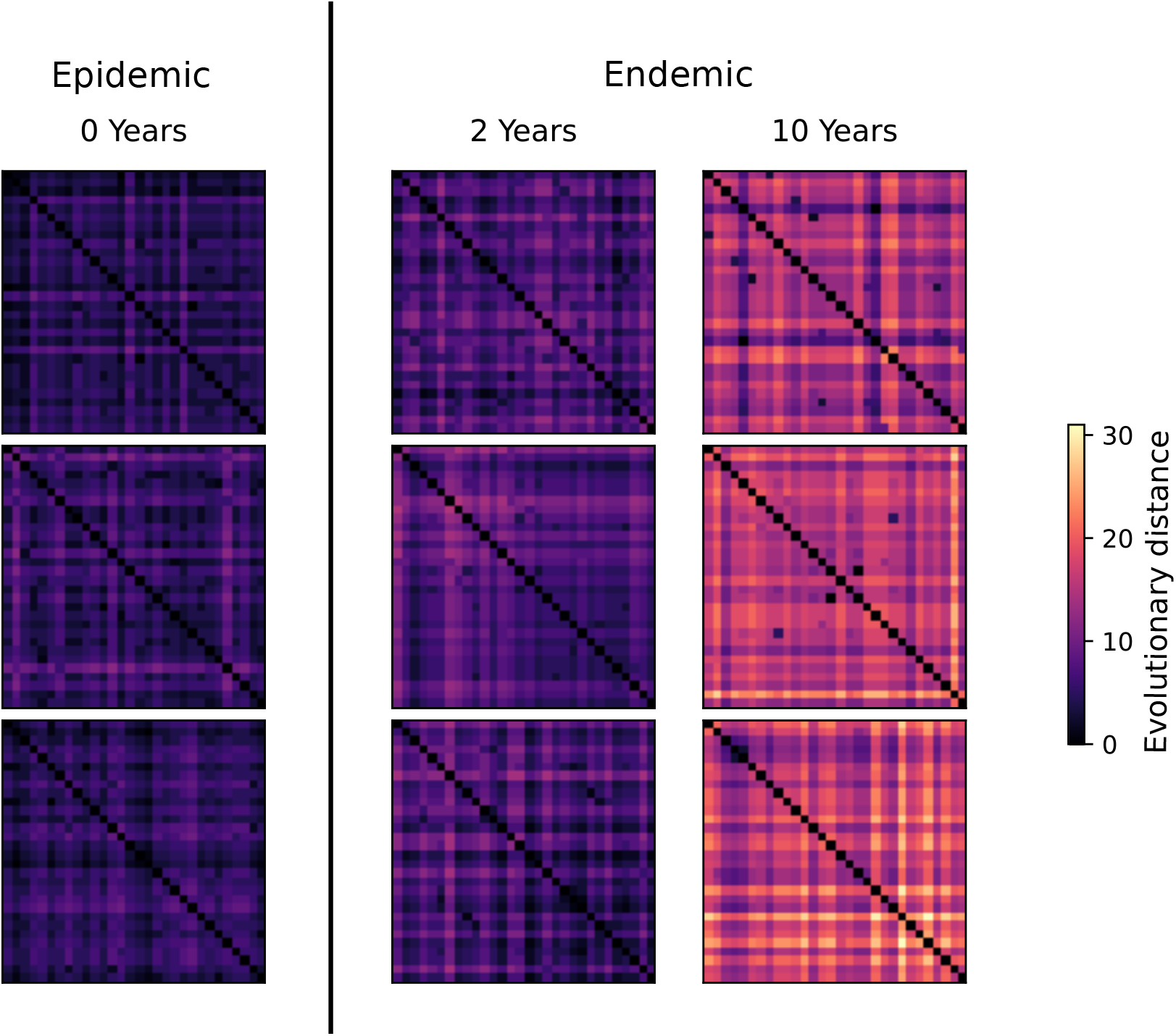
A collection of example images used for training from each training year. Each pictured training example was generated with *R*_0_ *>* 4. Three unsorted example images are shown for each sampling time (Fig 1A).

### The Convolutional Neural Network (CNN) Model

We developed a deep learning model using a Convolutional Neural Network (CNN) to solve a classification task predicting the label (0, 2, or 10 years) for a given pairwise distance matrix. The pairwise distance matrix is similar to a grayscale image (Fig 2). Thus, in the context of machine learning models, we also refer to a pairwise distance matrix as an image.

We constructed a CNN using Tensorflow [25] with a sequential architecture comprised of 2D convolutions, batch normalization, ReLu activations, and spatial maximum pooling. The layer structure is described in table 1. The inputs to the first and third dense layers were regularized using dropout with probability *p* = 0.25. We generated five variants of this model architecture to accept different input square matrices with side lengths of 15, 20, 30, 40, or 50 (corresponding to the number of sampled individuals in the pairwise distance matrix). We refer to this parameter as the input shape or the window size interchangeably. The number and size of all convolution kernels, size of max-pool filters, and dense layer output neurons were held constant across all models. Model architecture specifications are reported in table 1. Five batches of synthetic training data of 60,000 pairwise distance matrices were generated for each number of sampled individuals (for a total of 25 training sets), each with 20, 000 examples for each label. Each dataset was used independently to train a single neural network. Each neural network was trained within under an hour using two P100 GPUs with two Power8NVL CPUs. Five validation sets of 30,000 pairwise distance matrices (10,000 matrices per label) was used to compute and compare model performance, one set for each input size. Each model was trained with a batch size of 64 images using the Adam optimizer [26] for 50 epochs with a learning rate of 10^*−*4^. The first and second dense layers wer regularized with an *f*_2_ penalty with weights of 0.05 and 0.01 respectively. Models were evaluated independently at training time and as an ensemble in deployment using a majority voting system.

**Table 1.** The layer connectivity proceeding from the Input layer (top) down to the output layer. Our representation of image data within the convolutional stack is *channels last*. The batch size is implicitly 1 for each image and is dropped. If *k >* 20, an additional max-pool operation is performed before layer 10. No bias weight is used in the final dense layer.

| Layer index | Layer type | Output shape (per image) |
| --- | --- | --- |
| 0 | Input | $(k, k, 1)$ |
| 1 | Convolution 2D | $(k, k, 32)$ |
| 2 | Batch Normalization | $(k, k, 32)$ |
| 3 | Convolution 2D | $(k - 3, k - 3, 32)$ |
| 4 | ReLU Activation | $(k - 3, k - 3, 32)$ |
| 5 | Convolution 2D | $k - 5, k - 5, 32)$ |
| 6 | Max-Pool | $(\lceil \frac{k-5}{2} \rceil, \lceil \frac{k-5}{2} \rceil, 32)$ |
| 7 | Batch Normalization | $(\lceil \frac{k-5}{2} \rceil, \lceil \frac{k-5}{2} \rceil, 32)$ |
| 8 | ReLU Activation | $(\lceil \frac{k-5}{2} \rceil, \lceil \frac{k-5}{2} \rceil, 32)$ |
| 9 | Convolution 2D | $(\lceil \frac{k-5}{2} \rceil - 2, \lceil \frac{k-5}{2} \rceil - 2, 32)$ |
| 10 | Flatten | $((\lceil \frac{k-5}{2} \rceil - 2)^2 \cdot 32, 1)$ |
| 11 | Dropout | $((\lceil \frac{k-5}{2} \rceil - 2)^2 \cdot 32, 1)$ |
| 12 | Dense | $(64, 1)$ |
| 13 | Dense | $(64, 1)$ |
| 14 | Dropout | $(64, 1)$ |
| 15 | Dense | $(3, 1)$ |
| 16 | Softmax | $(3, 1)$ |

### Robustness of Model Prediction against Reordering

We tested the robustness of model predictions against reordering of elements of an image using a permutation test. We asked the question *what is the probability that randomly reordering k individuals represented in a pairwise distance matrix results in a different model prediction?*

To generate reordered images, we first selected *k* individuals to be re-ordered (combinations) and then reordered the *k* elements (permutations). To compute an average probability for a model, we computed all possible pairwise reorderings for 1000 randomly selected images. Each image generates 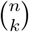 possible variations to consider. A subset of 1000 images were sampled from each validation set are used to compute the probability that random reordering results in the same model prediction, i.e. the measure for robustness. We considered both average accuracy and performance conditioned on initial model prediction (S2 Table).

### The Sliding-Window Approach

To handle a large number of sequences, we employed a sliding-window approach. We first constructed a pairwise distance matrix *D* of size *m* × *m* for sequences in a large database. We then used a window of size *n* × *n* and slided this window from one diagonal to the other diagonal of *D*. This method evaluates one *n* × *n* principal submatrix block in the sliding window at a time by assigning a label to the submatrix using the CNN model. For each individual represented in the pairwise distance matrix, we collected a list of the labels provided by the CNN model for each block they are represented in. The most frequently identified label within the list of predictions was assigned to an individual.

To validate the performance of the sliding window approach, we generated pairwise distance matrices of dimension 600 × 600 using our stochastic simulator. For each of the pairwise distance matrix, we performed 10 simulations using the same parameters as with the training dataset for the CNN model. In each simulation 60 individuals are sampled at either 0, 2, or 10 years. We then joined these 10 clusters by connecting their MRCAs by assuming that the distance between MRCAs of the clusters were uniformly distributed between 1 and 3 months. The order of individuals in the pairwise distance matrix was then randomized. A total of 100 independent pairwise distance matrices were generated with this process to form a validation set. Accuracy was computed on a per-individual basis using the sliding-window method.

### Outbreak Identification from Real Data

Applying the sliding-window approach to real HIV sequences led to predictions of sequences belonging to either an ’epidemic’ or an ’endemic’ phase. The prediction of a sequence collected at an earlier time may change when newly sampled sequences are added to the database, and this change may indicate an epidemiological link between an older sequence and newly sampled sequences. Therefore, for the sequences labeled as ‘epidemic’, we further considered three possible, inferred epidemiological situations based on the time a sequence was sampled relative to the time of the most recent samples (e.g. the current year).

For sequences sampled within 2 years of the current year, they were labeled as part of an *active epidemic*. For sequences sampled more than 2 years before the current year, we categorized them according to whether they were in the same sliding window as (i.e. close to) any sequences sampled in the current year that are labeled as ‘epidemic’. If so, we labeled these sequences *reactivated epidemic*; otherwise, we labeled them *inactive epidemic. Reactivated epidemic* indicates a situation where the newly identified outbreak may be linked to previously sampled sequences (i.e. individuals) in the database. *Inactive epidemic* indicates that we predict that the sequence used to belong to an outbreak at the time when the sequence is added to the database; however, it may not be relevant to current outbreaks.

The overall procedures of model prediction on an expanding database is represented in Fig. 3.

**Fig 3.**
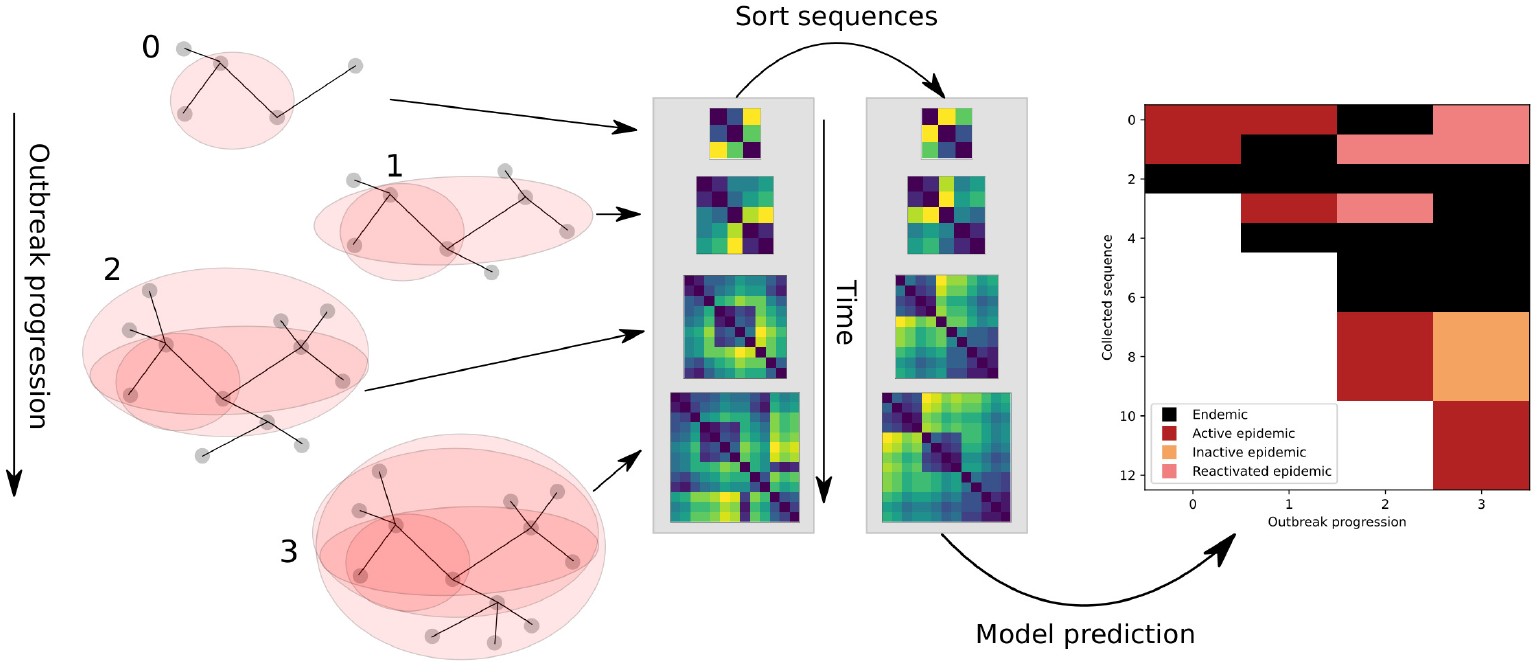
Cartoon representation of the evaluation of real data in real time. As the outbreak progresses and more sequences are collected (left), sequences are added to the database. Shaded areas indicate which host that are sampled at each time in a growing transmission network. At each time point (middle), we construct a pairwise distance representation of the sampled sequence data and apply a sorting algorithm. Collecting the predictions from each matrix gives a representation of the history and progression of an outbreak (right). Each step in the inference progression can add one or more new sequences from previous infections.

### HIV Sequence Data Used in the Study

To test our model performance in a realistic setting, where sequence data enters a public health database daily over long time, we compiled data on the HIV-1 CRF01 spread in Europe. The data was sorted on sampling year, and randomized within each year. This data contains a well-known dual outbreak of HIV among first Finnish, then Swedish, intravenous drug users [27]. The sequences from that dual outbreak thus become mixed with other sequences from Europe. We arranged this database to mimic the inflow of data from an outbreak that emerges in only part of the whole data as it enters the analysis stream, one new sequence at a time. In total, this data consisted of 277 sequences covering approximately 300 nt surrounding the HIV-1 env V3 region.

To determine the sensitivity of our method to the choice of evolutionary distance model used to calculate pairwise evolutionary distances, we analyzed HIV-1 subtype A6 sequence data from Former Soviet Union, Russia, and Ukraine (417 sequences covering approximately 400 nt surrounding the env V3 region. This dataset was sampled over a 20+ year range and features long distances where a correction factor may be significant. We computed pairwise distance matrices using a Hamming model (“Ham” which counts only mutations per site), JC69, K80, F81, and TN93 using Ape [28, 29]. Before evaluating the model, we subset the data by year to obtain pairwise distance matrices containing sequences sampled around the same time. Years with less than the minimum number of sequences needed to make a prediction were joined forward to the next year. The resulting pairwise distance matrices were evaluated using the sliding window method and accuracy values were calculated on a per-person basis.

To test scalability, we extracted HIV-1 RT (p51) gene sequences of at least 500 nt of any subtype or recombinant in the LANL HIV database (https://www.hiv.lanl.gov/content/sequence/HIV/mainpage.html). We then removed all sequences labelled as “problematic” in the LANL HIV database and further all that required gaps, as RT typically does not have many gaps, resulting in a 1,320 nt alignment of 271,868 sequences. To estimate the runtime of our pipeline, we subsequently resampled this alignment for 10^2^ − 10^5^ sequences and constructed the pairwise distance matrix using the TN93 evolutionary model. We also measured the time to sort each matrix using hierarchical clustering and the time to stride the larger matrix into smaller views and pass the data through the model.

All HIV datasets were aligned using MAFFT version 7 [30].

## Results

### The convolutional neural network (CNN) model classifies synthetic data with high accuracy

We developed a framework employing CNN models to identify active outbreaks from HIV-1 sequence data (see Methods for the workflow and details of the framework). We first generated synthetic datasets for training the CNN models using our forward stochastic simulator of HIV transmission for training and testing of the CNN model. We varied the reproductive number, *R*_0_, between 1.5 and 5 in the simulator. Samples were taken at three different time points, i.e., during an ongoing outbreak, at year 2 after the infected population growth had ended, or at year 10 after the population growth had ended (Fig 1A and Methods), corresponding to label 0, 1 and 2, respectively, in the CNN model. At each time point, a total of 15, 20, 30, 40, or 50 samples are taken. The sample size here is used as the sliding window size later in the general framework. For each sample size, we repeated the stochastic simulation to generate 300,000 pair-wise distance matrices. The memory requirements to store the full training set for the larger sample sizes contiguously in GPU memory exceeds the capacity of many consumer-grade laptops and desktops. Therefore, we split them into five subsets of matrices with equal size, i.e., 60,000 labeled pairwise distance matrices.

We then trained a CNN model on each of the five window size subsets of data to correctly predict the labels (i.e. label 0, 1 and 2) of the pair-wise matrices. Similar model performance were achieved across the five subsets (Fig. 4A). We then joined all the trained networks in a simple bagging classifier with a voting decision method. This led to a classifier with a higher accuracy than the mean accuracy of the five networks used to assemble the larger classifier model (Fig. 4B). We thus used the voting classifier for predictions below. Overall, the accuracy of the CNN model with the voting decision method ranged between 79% and 87%, and the accuracy improved with increased sample size (i.e., larger windows perform better; Fig. 4).

**Fig 4.**
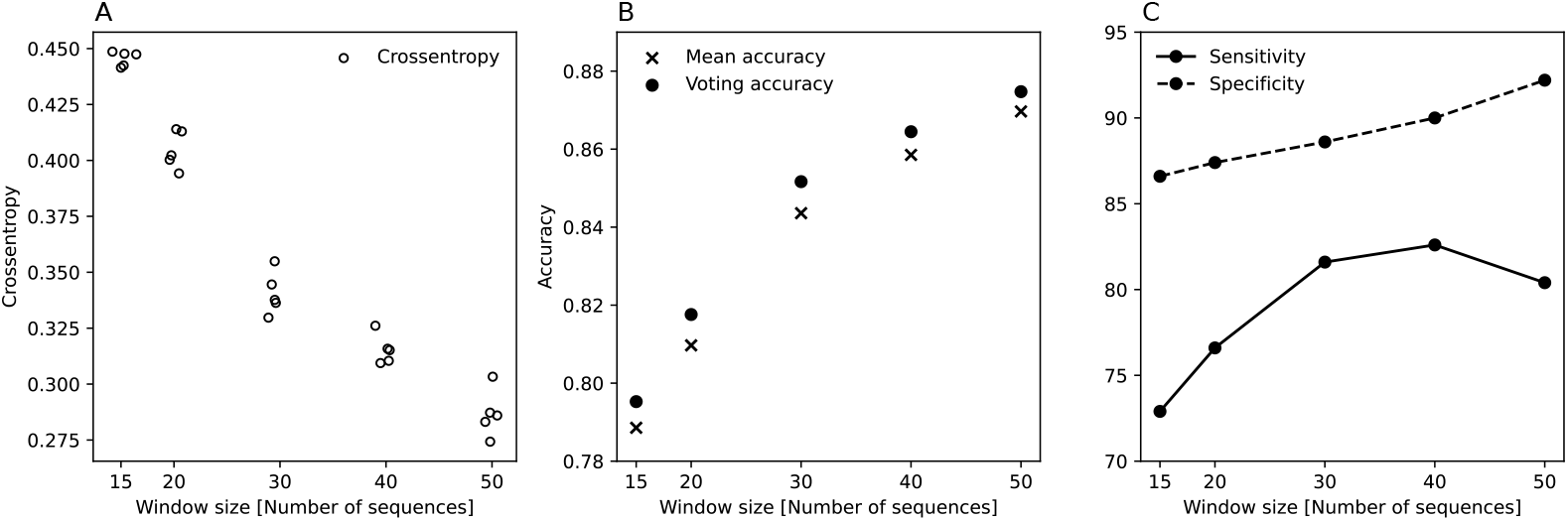
(A) Crossentropy of each trained neural network on the test sets, a statistical measure of the average probability assigned to the incorrect labels by each model. Performance of networks trained on each subset of data are similar to networks trained on other subsets. (B) Accuracy of trained models on the same test sets. Constructing a voting classifier model from individual neural networks improves accuracy beyond using a single trained network by 0.5−2%. (C) Sensitivity and specificity of the voting models on the binary problem of identifying epidemic from endemic samples. Specificity (true negative rate) improves with a larger window, but specificity does not always improve with a larger window.

From a public health perspective, we are interested in identifying sequences in a outbreak (i.e. label 0 in our CNN model) among sequences collected from an endemic (non-outbreak; label 1 and 2 in our CNN model) phase. We thus examined the specificity and sensitivity of our model in terms of identifying the outbreak (i.e. correctly predicting label 0). We classified label 0 as epidemic, and label 1 and 2 as ’endemic’, and found that at all window sizes, the specificity (of identifying ’epidemic’ sequences) exceeds 85% and the sensitivity exceeds 70%. Specificity increases with the addition of data, while sensitivity for our method increases initially, but appears to saturate at window sizes of 40 and 50.

An important feature of our framework is that it does not rely on phylogenetic tree construction; instead, it used a clustering algorithm to group closely related sequences together. While phylogenetic tree construction is more reliable to specify evolutionary relationships (and thus the genealogical relationships) of the sequences, it takes a longer time to perform than clustering algorithms. Here we show that precise evolutionary relationships is not necessary for our CNN model to make correct predictions as long as closely related sequences are grouped together (through clustering algorithms). To this end, we permuted two randomly chosen sequences in a matrix and calculate the probability that the model gives the correct prediction (S2 Table). Note that reordering of the sequences could lead to a different image (pairwise-distance matrix) on which the CNN model makes its prediction. Overall, a random reordering of two sequences preserves a correct prediction with a probability between 80% and 89%. This suggests that our method is robust to the specific ordering of the sequences presented to the model.

### The convolutional neural network based model performs better when the outbreak is more explosive

The basic reproductive number *R*_0_ is one of the key parameters characterizing an outbreak. We analyzed the performance of our model in relation to the *R*_0_ parameter used during simulation. Matrices in the training set were partitioned by label (’epidemic’ or ’endemic’), and then binned according to the value of *R*_0_ into 250 uniformly sized bins. Figure 5 shows the average accuracy for each such bin. Our model performed well at identifying sequences taken from the endemic phases. In terms of identifying sequences sampled from an epidemic, the model accuracy exceeded 60%, when *R*_0_ *>* 3. Epidemics with *R*_0_ less than 3 were difficult to detect. However, increasing the sample size increased the probability of identifying an outbreak notably. We further evaluated how the maximum population size in the simulator impacts the accuracy of model prediction (S1 Fig). In general, the model performance is less impacted by the choice of this parameter.

**Fig 5.**
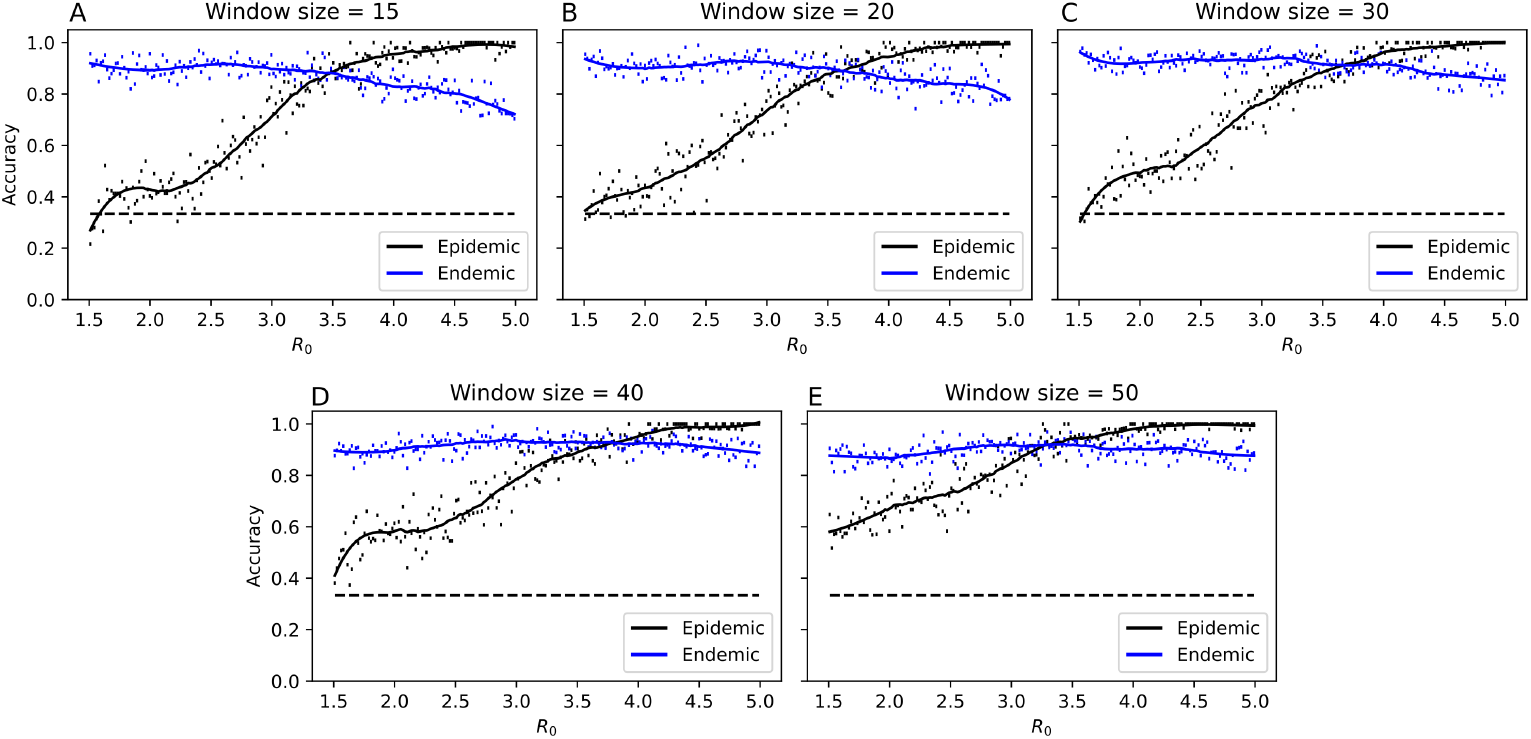
Association between *R*_0_ parameter and per-image model performance by epidemic vs endemic training label. The *R*_0_ axis is subdivided into 200 subintervals. Accuracy is calculated over each subinterval, a trend line is fitted using a Savitzky-Golay filter of degree 3 with a window size of 51. A black dashed line indicates the probability of predicting an outbreak using only the occurrence rate in the training set, independent of *R*_0_. Outbreaks with small *R*_0_ are more difficult to detect than with larger *R*_0_.

### The sliding window approach identifies subsets of sequences taken from outbreaks

To identify the subsets of sequences that were sampled from an outbreak in a database of many sequences, we used a sliding window approach. In this approach, we first constructed a pairwise distance matrix for all the sequences in the database, then sort the matrix so that closely related sequences are grouped together. We then employed the sliding window to assign predictions to groups of individuals in the matrix (see Methods for details).

We considered three different sorting methods: 1) randomly ordered, 2) hierarchical clustering (Ward criterion), and 3) hierarchical clustering (Ward criterion) with optimal leaf ordering refinement. To test the performance of the three methods, we generated 100 pairwise matrices (images) of size 600 by 600. Each of the 600 by 600 matrix was generated by joining 10 simulated matrices using the stochastic simulator with outbreak parameters (*R*_0_ and simulation length) randomly sampled (see Methods for details). In total, we evaluated 60,000 predictions to compute the accuracy of these model predictions. Of the three tested sorting methods, applying hierarchical clustering gave the best accuracy (see S3 Fig). Thus, hereafter, we applied this sorting method to all large matrices. This approach also achieved high sensitivity and specificity in detecting outbreaks (figure 6).

**Fig 6.**
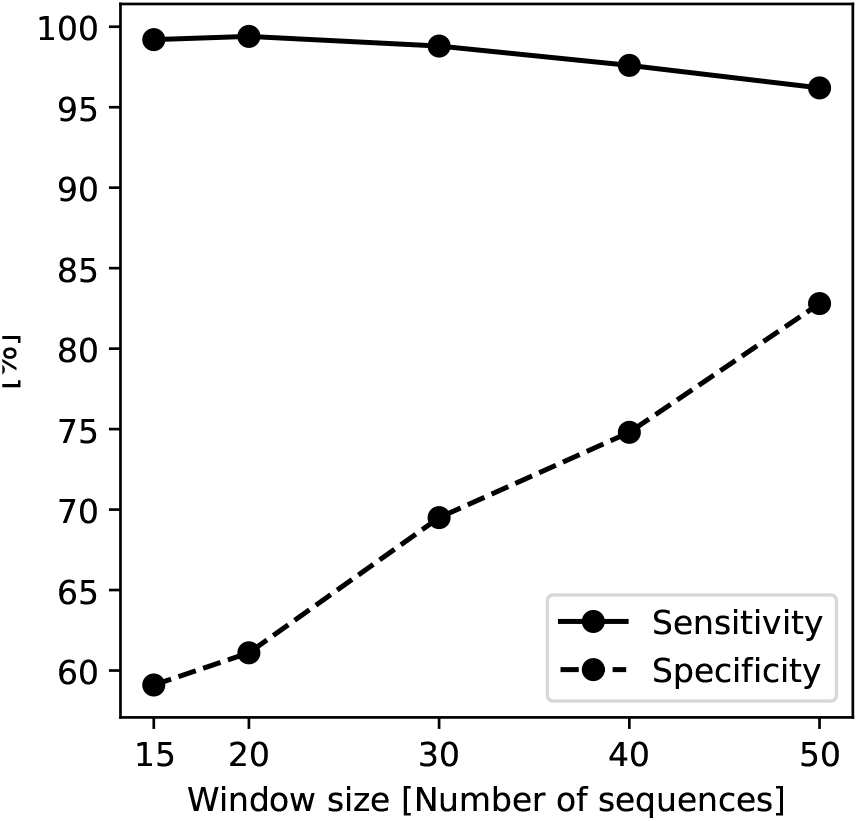
Sensitivity and specificity on the multiple-cluster test set. Model performance is computed using the same 60, 000 infections. Sensitivity is high but decreases as the window size increases, while the specificity of the method increases as the window size increases.

We also tested the robustness of model predictions using different evolutionary models to compute pairwise evolutionary distances from real HIV-1 sequence data. We used a meta-dataset of HIV-1 sequences from the former Soviet Union [31] (see Methods). Overall, for models using all window sizes (15, 20, 30, 40, 50), there is minimal pairwise variation between the predictions using different evolutionary models (S2 Fig).

### Validating model using historical HIV-1 data

To validate our model against real datasets and illustrate the application of a real outbreak identification in real time, we collected a dataset containing a well-documented dual outbreak among first Finnish and then Swedish IDUs [27]. To add complexity, we added an assortment of other European HIV-1 infections from the time period around the Finnish-Swedish outbreak to test our model’s ability to detect an outbreak in an endemic backdrop (Fig 7). An initial subset of 15 sequences is required to make a first prediction (‘circled 1’ in Fig 7). These initial 15 sequences were classified as endemic, which agreed with how they were sampled; coming from different European countries (including Sweden), not as part of any known outbreak. Note that most of these initially classified endemic sequences stayed endemic throughout the growth of the database.

**Fig 7.**
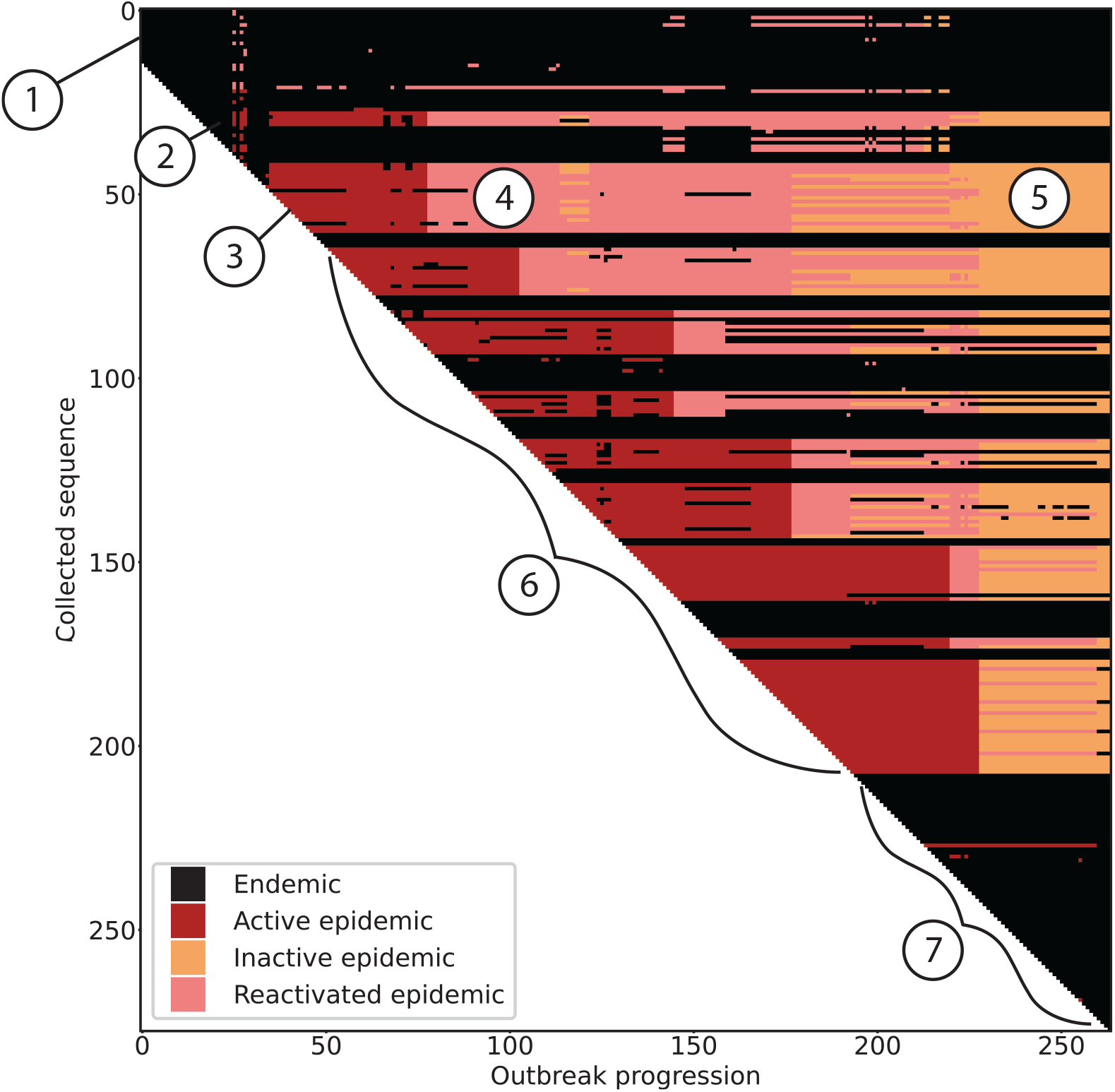
Real-time detection of a HIV-1 CRF01 outbreak. As sequences are added to a growing database (from top to bottom on the y-axis) and time progresses (x-axis), one distance matrix is analyzed at each sequence addition and an updated classification of all current sequences is added as a slice along the x-axis. Our model classifies each infection as belonging to an outbreak (epidemic) or as endemic. The epidemic label is refined into active, inactive, or reactivated based on the time since detection. Here, the transition cutoff from active epidemic to inactive/reactivated epidemic is 2 years. Circled numbers indicate events of interest, discussed in the main text.

After the initial 15 sequences, the database grows one sequence at a time, and we calculated a new distance matrix for all sequences up to that point, applied sorting, and performed predictions using the sliding window as outlined in Figure 1. The sequences were re-classified in each matrix step, and the result added as a slice along the x-axis. At ‘circled 2’, additional sequences were added as endemic, but shortly thereafter, when sequences were added at ‘circled 3’, their classification changed to ‘active epidemic’ as new sequences inform about an ongoing outbreak. This correctly identified the start of the Finnish IDU outbreak in 1999. Sequences that joined the database at ‘circled 3’ and were classified as active epidemic, later became classified as ‘reactivated epidemic’ at ‘circled 4’ as they were about to become ‘inactive epidemic’ but new active epidemic sequences were added, instantly reactivating them. At ‘circled 5’, these (and many other) sequences eventually became inactive epidemic state. During ‘circled 6’, first, the Finnish outbreak developed as shown in red; interspersed with unrelated sequences from patients in Denmark and the UK added correctly in black; then, while Finnish sequences kept being added in red, some Swedish IDU sequences were added in black around year 2001, these were correctly identifying pre-outbreak slow spread in Sweden [27], again with various sequences from other European countries that were not part of any known outbreak also added in black. Eventually, in 2004, the well-documented Swedish IDU outbreak started, accurately identified by our model. At the end of the database growth, around ‘circled 7’, the final 69 sequence additions (except one) joined as endemic state (black). This involved both Finnish and Swedish post-outbreak slower spreading HIV in the IDU group, as well as other European sequences from the same time period. At the final slice in the figure, all sequences were classified as either ‘endemic’ or ‘inactive epidemic’, correctly indicating no new outbreak in these data at that time. Overall, our model correctly identified the epidemiological dynamics of the Finnish-Swedish IDU outbreaks among other European HIV-1 CRF01 sequences during the same time period as the outbreaks happened. Importantly, we detected both outbreaks early on, before they were over, implying that had this method been applied in real-time as data became available, we would have been able to intervene and possibly prevent the extent of the outbreaks.

### Our approach efficiently handles over 100,000 sequences

Finally, we evaluated the ability of our model to perform under increasing quantities of sequences. We downloaded 271,868 aligned HIV-1 pol sequences from the LANL HIV sequence database and subsampled sets ranging from 10^2^ sequences to 10^5^ sequences (see Methods). We measured the real time it took to construct a large pairwise distance matrix from the sample using Ape [28], the time required to sort the matrix using hierarchical clustering [32], and the time required slide the window down the sorted matrix diagonal and pass the data through the model to identify sequences belonging to an epidemic. Measured times were averaged at each data set size then log transformed and plotted in figure 8. A set of 100, 000 sequences were analyzed with this pipeline in under 70 minutes.

**Fig 8.**
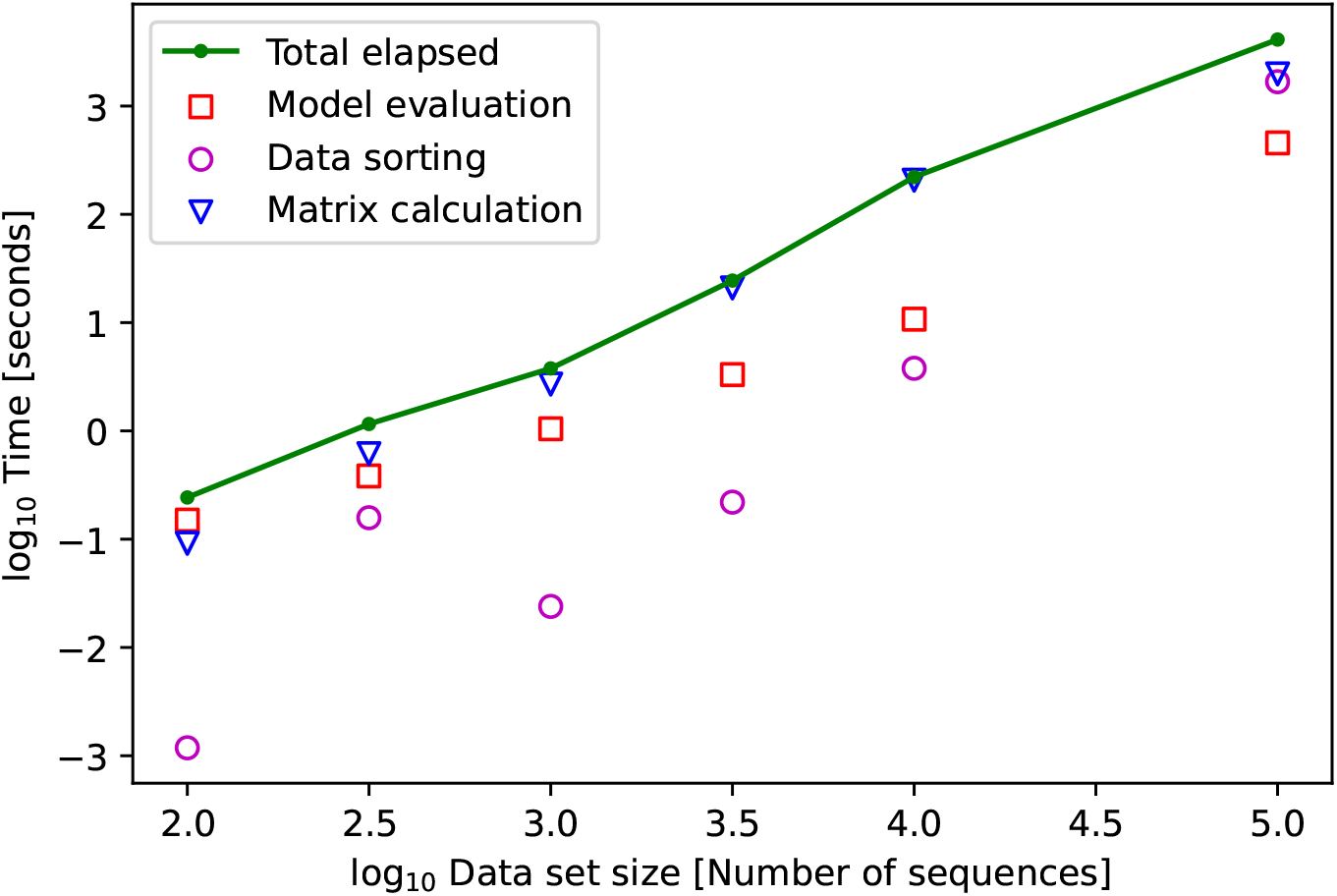
Decomposition of measured total method runtime into the primary components of distance matrix construction, distance matrix sorting, and model evaluation. Measured time performance averaged then log transformed. Less than 100 sequences can be analyzed in under 1 second.

In addition to the empirical results, we consider the computational complexity of these operations as the number of sequences in the dataset, *n*, increases. The sliding window method for annotating large matrices has *O*(*n*) time complexity as the number of model evaluations and the complexity of such evaluations grows linearly with the number of infections presented. Constructing the pairwise distance matrix has *O*(*n*^2^) time complexity, and hierarchical clustering implemented with nearest neighbor chaining is performed with *O*(*n*^2^) time complexity [18] giving a time complexity approximately *O*(*n*^2^) for the full pipeline.

## Discussion

In this work, we developed a deep learning approach to effectively and efficiently handle large quantities of viral sequences to identify subsets of sequences that were taken from an HIV-1 active outbreak. In contrast to phylogenetic approaches, our approach constructs and directly analyzes pairwise distance matrices from viral sequences. The key idea is to treat these matrices as images and then train and use convolutional neural network (CNN) models to identify subsets of sequences which form matrices that bear the signature of an outbreak. Using historical HIV-1 genetic sequence data with known epidemiological history, we showed that our approach correctly identified subsets of sequences sampled from outbreaks. Furthermore, we showed that our approach is scalable and can process hundreds of thousands of sequences within hours.

The key to the low-computational cost of our approach is analyzing a pairwise distance matrix derived from sequences in a database, instead of constructing phylogenetic trees. While computing large trees from the full sequences is possible using fast, but approximate, methods, subsequent interpretation is ad hoc, often subjective, and requires careful analysis by experts. Other, more advanced methods that partially incorporate interpretation and integrating across many plausible trees, such as elegant Bayesian methods [11], are instead even slower and unsuitable for very large data analyses. On the other hand, the dimension of the data encoded in a pairwise distance matrix is significantly less than that encoded in the full sequences. Importantly, it has been shown that this dimension reduction does not substantially impact on the information content with respect to evolutionary relationships between sequences [33]. By building the pairwise distance matrix, we concentrated the epidemiological signatures (outbreak or endemic) into a more compact structure, i.e. an image format, that can be analyzed very efficiently using deep learning. Given the trained system, the computational speed and complexity of our approach is currently limited by the quadratic complexity (in the number of sequences considered) in building the pairwise distance matrix and secondarily by the data sorting and model evaluation.

The accuracy of the model prediction is ensured by applying a deep learning method based on CNN models for image classification. The model was trained using a large set of synthetic data generated from a previously validated HIV-1 transmission simulator [20]. Our deep learning model performed well when *R*_0_ is high (*R*_0_ *>* 3). Large *R*_0_ characterizes fast-growing outbreaks that we would like to identify early and with high confidence. When *R*_0_ is small, the outbreak grows significantly slower and is difficult for our model to detect. More sophisticated CNN models than proposed here may be used in future improvements to the ability of the model to detect outbreaks. However, another factor that may lead to poor model performance is lack of signal in the data. For example, it is not clear if a reduction of *R*_0_ from 1.5 3 down to 1 for only 2 years, when the mean time to detection is about 2 years, induces strong signals of an outbreak in viral genetic data. Note that, increasing the sample size from 15 sequences to 30 or 50 sequences does not give a substantial increase in performance for low *R*_0_ S1 Fig. This further suggests lack of signal for detection.

Our approach is limited by the use of synthetic data obtained from a simulator to train the model. Synthetic data may lack certain subtleties present in real data. Thus, we validated our final model using real HIV-1 sequence data. We showed that detecting a known dual outbreak among injecting drug users first in Finland and then in Sweden, intermingled by random data from the same subtype in Europe, accurately identified the outbreak in a data stream where HIV sequences were provided over the time the outbreaks occurred. Encouragingly, we detected the outbreaks early, before they were over, as well as the end of the outbreaks. Accurately identifying a beginning outbreak in the background of other data and knowing when it’s over are both crucial pieces of information for a public health authority.

Our approach has potential to effectively deal with recombination when analyzing sequences. Standard phylogenetic methods assume no recombination, which HIV clearly violates. Pairwise distances, on the other hand, are accurate whether recombination took place or not. Here, our deep learning model is trained to classify sets of evolutionary distances in very large data collections with no requirement to perform manual or automated cluster separation and examine only regions of high density or low genetic divergence [34]. Our technique can identify sequences likely to be part of an epidemic without the need for a strict evolutionary distance cutoff and avoids the need to a priori give a strict definition of a cluster.

Overall, our deep learning approach is capable of processing tens of thousands of sequences, and is not dependent on a specific choice of evolutionary distance metric. While we developed this computational framework for HIV-1, with adjustments in the evolutionary rate and training simulations, it could be applied to analyze other pathogens for online, rapid response to developing outbreaks and rising of novel variants of concerns, e.g. for SARS-CoV-2.

## Data availability

The source code used to produce the results and analyses presented in this manuscript are available from: https://github.com/MolEvolEpid/MachineLearningModelforHIVOutbreaks

## Author contributions

Conceptualization: R.K.; Methodology: T.L. and R.K.; Formal Analysis: M.D.K. and R.K.; Funding Acquisition: T.L. and R.K.; Writing: M.D.K., T.L. and R.K.

## Supporting information

**S1 Table. Number of parameters specified for each neural network and parameters used in the dense layers**.

**S1 Fig. Analysis of accuracy as a function of both population size (***N*_*pop*_**) and** *R*_0_. Each dimension is binned into 30 subintervals. Accuracy is calculated on 10, 000 sample images for each combination of outbreak status and window size.

**S2 Table. The probability of reordering two individuals within the matrix is a easure of model performance against reordering**. A subset of 1000 matrices are sampled from the test set and all pairwise reorderings are constructed, and then evaluated with the model. We report the overall accuracy on the reorderings and the accuracy conditioned on the correctness of the initial prediction.

**S2 Fig. Similarity of model predictions on Russian dataset when presented by year**. Models with smaller window sizes give similar predictions, an advantage over models with larger window sizes.

**S3 Fig. Performance of the sliding window algorithm with different clustering methods applied to the full data set**. Method HC denotes hierarchical clustering, OLO denotes HC with optimal leaf ordering refinement, None denotes no sorting is applied. Accuracy is computed using the three labels used during training.

