## Supplementary Material for "A deep learning approach to real-time HIV outbreak detection using genetic data"

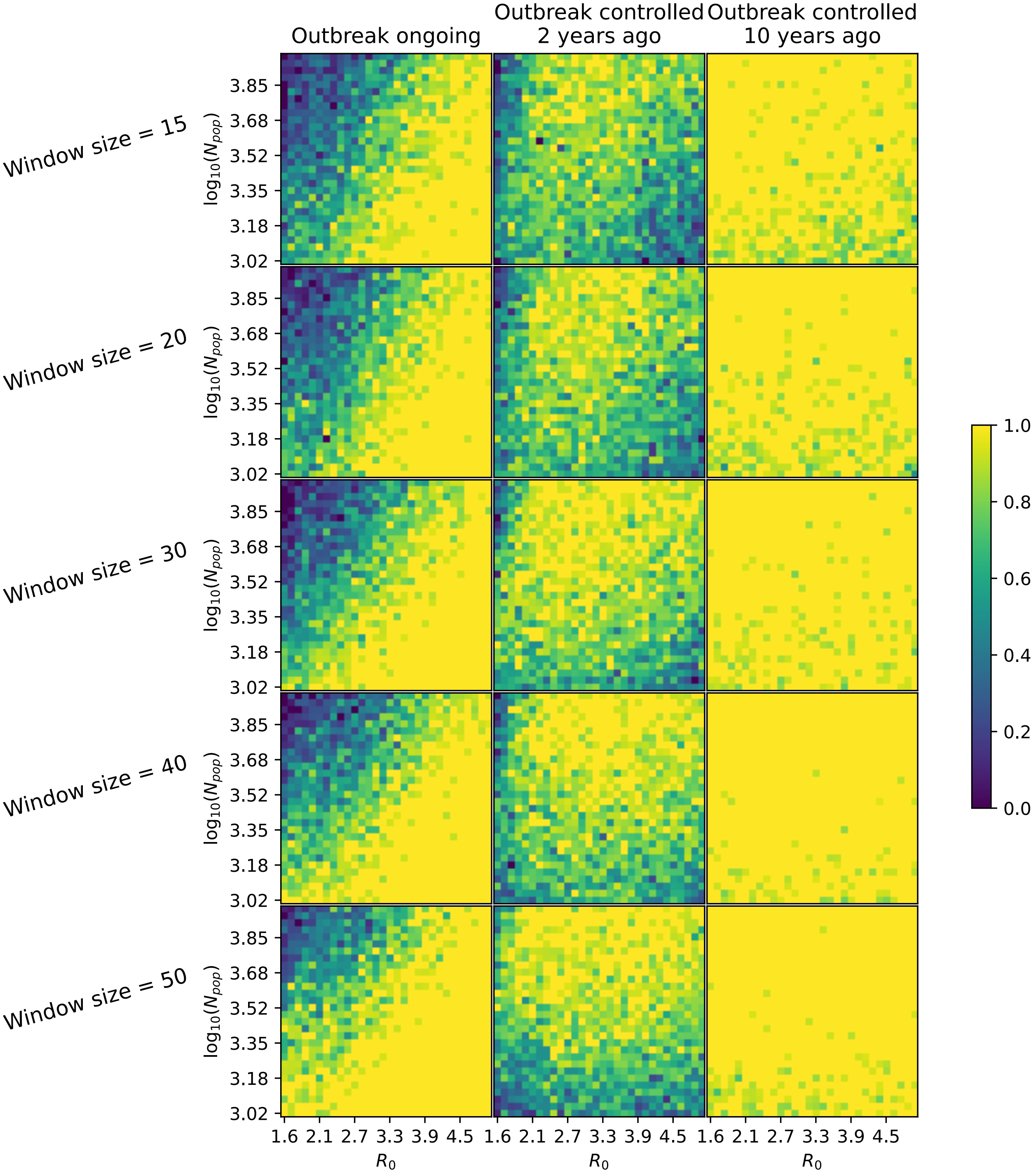


Figure S1: Analysis of accuracy as a function of both population size and $R_0$. Each dimension is binned into 30 subintervals. Accuracy is calculated on 10,000 sample images for each combination of outbreak status and window size.

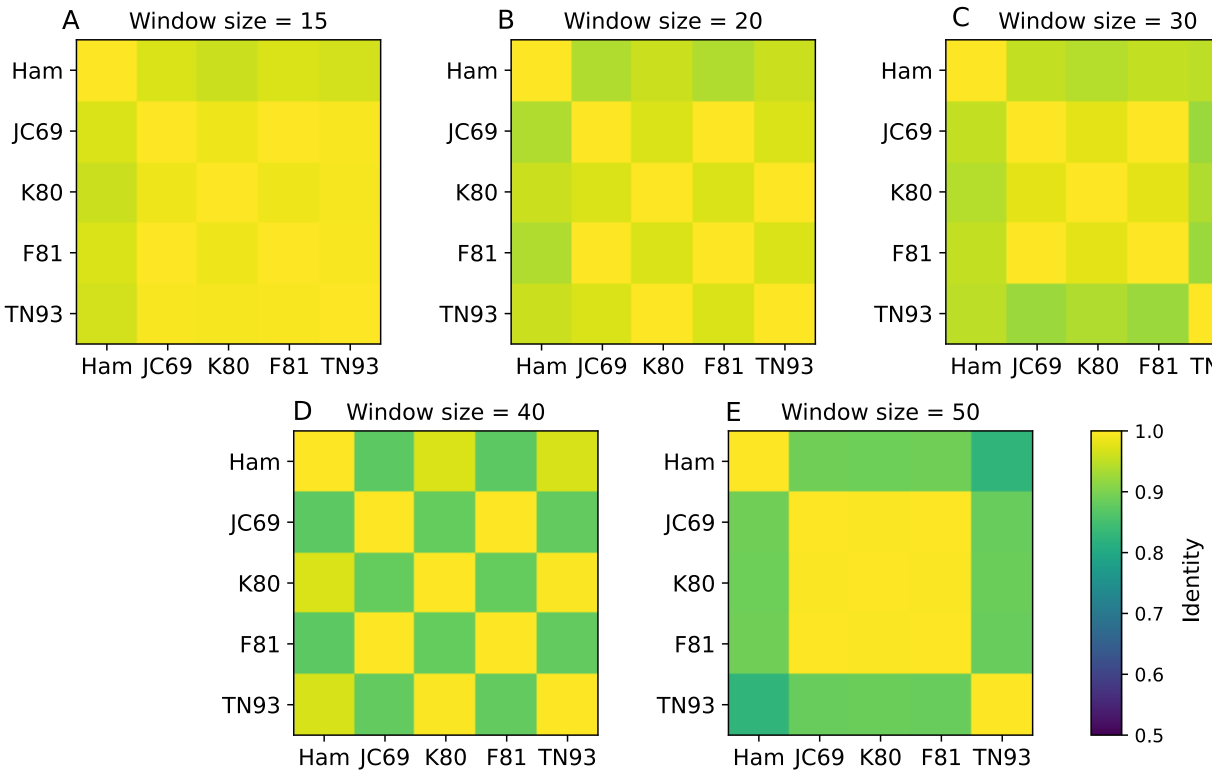


Figure S2: Similarity of model predictions on Russian dataset when presented by year. Models with smaller window sizes give similar predictions, an advantage over models with larger window sizes.

Table S1: **The number of trainable parameters in models increases as the number of samples used to generate each input matrix increases.**

| Sample Size ($k$) | Trainable Parameters | Trainable Parameters in Dense Layers |
|---|---:|---:|
| 15 | 58,432 | 22,848 |
| 20 | 91,200 | 55,616 |
| 30 | 91,200 | 55,616 |
| 40 | 140,352 | 104,768 |
| 50 | 244,800 | 209,216 |

Table S2: **The probability of reordering two individuals within the matrix is a measure of model performance against reordering.** A subset of 1000 matrices are sampled from the test set and all pairwise reorderings are constructed, and then evaluated with the model. We report the overall accuracy on the reorderings and the accuracy conditioned on the correctness of the initial prediction.

| Window size | Accuracy | Accuracy given initial prediction correct | Accuracy given initial prediction incorrect |
|---|---:|:---:|:---:|
| 15 | 0.808457 | 0.974737 | 0.117624 |
| 20 | 0.817047 | 0.979186 | 0.093184 |
| 30 | 0.869834 | 0.985382 | 0.103343 |
| 40 | 0.882845 | 0.992225 | 0.080726 |
| 50 | 0.889078 | 0.992451 | 0.108920 |

Calculated probability that a reordering of any two individuals gives the correct prediction on 1000 matrices sampled from the validation set used during training.

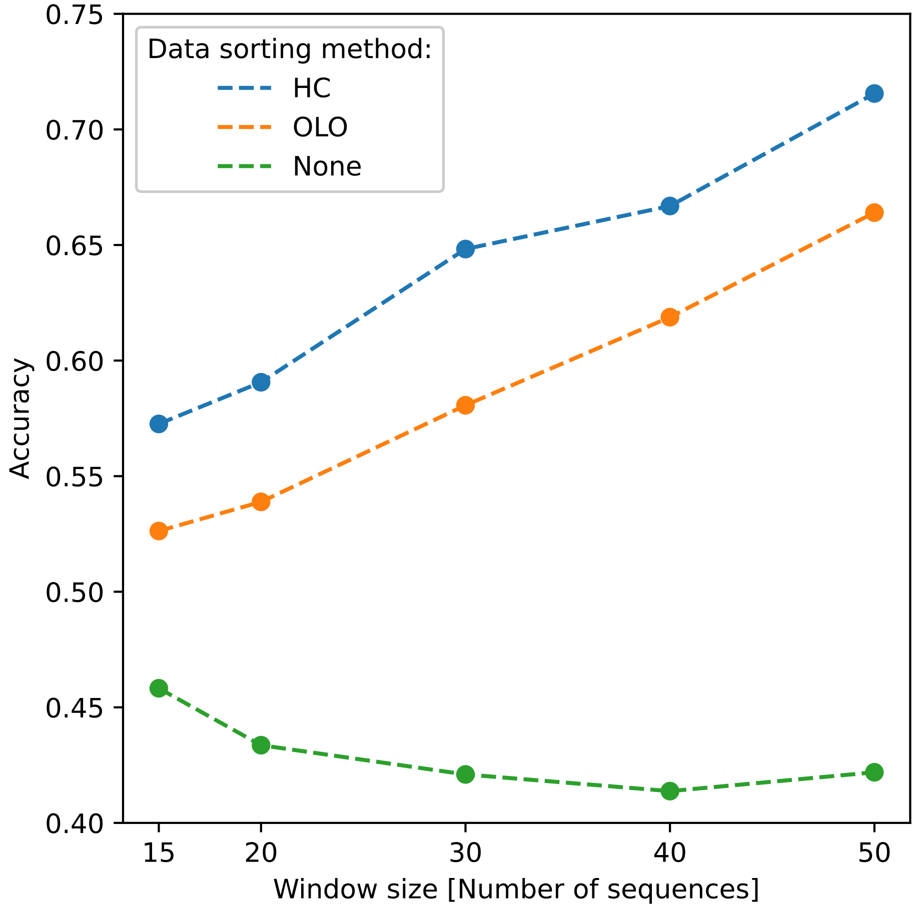


Figure S3: **Performance of the sliding window algorithm with different clustering methods applied to the full data set.** Method HC denotes hierarchical clustering, OLO denotes HC with optimal leaf ordering refinement, None denotes no sorting is applied. Accuracy is computed using the three labels used during training.
